# Lipid composition-driven sorting of ABHD5 between monolayer and bilayer surfaces

**DOI:** 10.64898/2026.08.20.744708

**Authors:** Shahnaz Parveen, Arvin Nazari, James Stebelton, Kahlid Ali, Matthew Sanders, Amit Kumar, Katherine Truex, Yu-Ming M. Huang, James G. Granneman, Christopher V. Kelly

## Abstract

Lipid droplets (LDs) are ubiquitous organelles that store neutral lipids and serve as central regulators of lipid homeostasis. Their structure includes a hydrophobic core of triacylglycerols and sterol esters surrounded by a phospholipid monolayer. This organization creates biophysical properties that guide selective protein recruitment. Among LD-associated proteins, α/β-hydrolase domain-containing protein 5 (ABHD5, also known as CGI-58) is a key regulator of lipolysis and broader lipid metabolism, yet the mechanisms guiding its distribution between endoplasmic reticulum (ER) bilayers and LD monolayers remain poorly understood. Because proper membrane association of ABHD5 is essential for activating PNPLA family lipases, identifying the determinants of its membrane selectivity is critical for understanding LD function in health and disease. In this study, we examined ABHD5 binding and sorting behavior using model membrane systems composed of giant unilamellar vesicles (GUVs) and droplet-embedded vesicles (DEVs) incorporating defined phospholipid and neutral lipid compositions. By integrating experimental assays with computational modeling, we quantified how ABHD5 partitions between bilayer membranes mimicking the ER and monolayer surfaces mimicking LDs. Systematic variation of membrane composition and physical properties allowed us to assess how packing defects and neutral lipid content shape ABHD5 localization. Our findings reveal the biophysical features that favor ABHD5 association with LD-like monolayers and provide new mechanistic insight into how cells target regulatory proteins to distinct membrane environments to control lipid metabolism.

## Introduction

Lipid droplets (LDs) are prevalent cellular organelles that store neutral lipids such as triacylglycerols and sterol esters, acting as dynamic energy reservoirs and critical hubs for lipid homeostasis (1, 2). Lipid droplets have a unique structure with a hydrophobic neutral lipid core surrounded by a phospholipid monolayer. This structure distinguishes them from typical bilayer organelles and allows selective recruitment of proteins that control lipid storage, mobilization, and signaling (3–5). Proteins partition to the LDs from the cytosol (CYTOLD), from the endoplasmic reticulum (ER) (ERTOLD), or via association with other proteins (INTOLD) (5, 6). Dysregulation of LDs is implicated in a spectrum of metabolic disorders, including obesity, fatty liver disease, and other lipid storage diseases (7–10). In this context, understanding how lipids govern protein partitioning is crucial for explaining how cells regulate lipid metabolism at the organelle level.

Protein targeting between organelle surfaces is known to be dependent on lipid composition (11). Variations in phospholipid species and neutral lipid content can modulate how proteins distribute between distinct lipid environments via specific lipid binding pockets, associations with charged moieties, curvature sensing, or non-specific sensing of the phospholipid biophysical properties (12–14). Previous studies indicate that LD monolayers exhibit a higher density of phospholipid packing defects than bilayer membranes, a property that is thought to be key for the recruitment of LD-associated proteins (15, 16). The specific lipid species, shape, and tension of the phospholipids largely determine packing defects (11–13, 16).

Among LD-associated proteins, α/β-hydrolase domain-containing protein 5 (ABHD5, also known as CGI-58) is a key regulator of lipid metabolism. ABHD5 lacks intrinsic hydrolase activity but functions as a coactivator patatin-like phospholipase domain-containing protein (PNPLA) lipases on the ER and LDs. ABHD5 regulates PNPLA2-mediated lipolysis on LDs (10, 17), PNPLA1-mediated skin barrier function at the ER (18, 19), and PNPLA3-mediated hepatic lipid release at the ER (20, 21), in addition to Hepatitis C virus assembly processes (22) and tumor progression (23, 24) via unknown mechanisms. Genetic inactivation of ABHD5 causes Chanarin-Dorfman syndrome, characterized by widespread accumulation of neutral lipids and severe ichthyosis (25, 26).

ABHD5 associates with organelles through direct lipid engagement and via interactions with resident proteins, such as perilipins (PLINs). Early work showed that PLIN1/PLIN5 bind ABHD5 and can recruit/stabilize it at LDs, leading to the INTOLD model (27, 28). Later work demonstrated that ABHD5 can directly interact with PNPLA-family proteins and recruit PNPLAs to specific lipid surfaces, providing a PLIN-independent protein-partner mechanism (29–31). ABHD5 acts as a lipid-sensitive molecular sensor and transducer that responds to cellular and lipid cues, regulating PNPLA localization and activity on LD and ER surfaces to coordinate diverse lipid metabolic processes.

The binding of ABHD5 to phospholipid surfaces is mediated by the N-terminal region (residues 1-43) and a membrane insertion domain within the insertion segment (residues 180-230), which together facilitate engagement with the lipid surface (22–25). These regions contain hydrophobic and aromatic residues that contribute to lipid interactions and surface anchoring (32). We hypothesize that ABHD5 localization is influenced by the presence and extent of phospholipid packing defects, which are regions where the membrane hydrophobic core and its neutral lipid content become partially exposed to the aqueous cytoplasm (12, 15). However, how these lipid-dependent effects contribute to ABHD5 partitioning between phospholipid bilayers and LD monolayers remains unclear. A deeper understanding of ABHD5-lipid interactions is therefore important for elucidating principles governing its localization in cellular lipid metabolism. In particular, it remains unclear how ABHD5 senses or responds to the distinct biophysical properties of bilayer versus monolayer membranes such as curvature, lipid composition, and packing defect density to achieve organelle-specific localization during lipolysis.

In this work, we investigated how ABHD5 sorts between surfaces of varying phospholipid and neutral lipid composition. We used model membrane systems and combined experimental measurements with computational modeling to determine how ABHD5 partitions between phospholipid bilayers and monolayers by systematically varying lipid composition and following its localization. In sum, we identified the importance of neutral lipids in imparting phospholipid packing defects that recruit ABHD5.

## Material and Methods

### Materials

1,2-dioleoyl-sn-glycero-3-phosphocholine (DOPC), 1,2-dioleoyl-sn-glycero-3-phosphoethanolamine (DOPE), L-α-phosphatidylinositol (Soy PI), and cholest-5-en-3ß-yl heptadecanoate (Cholesteryl ester) were purchased from Avanti Polar Lipids (Alabaster, AL). Triolein and Diolein were purchased from Nu-check Prep (Elysian, MN). The fluorescent probe NR12S membrane polarity probe was obtained from either TOCRIS Bioscience (Bristol, UK) or Cytoskeleton Inc. (Denver, CO, USA); no differences in probe performance were observed between suppliers. 96-well glass bottom plates (1.5 H, uncoated) were purchased from Cellvis LLC (Mountain View, CA), and 24-well glass bottom plates (1.5 H, uncoated) were purchased from MatTek Corporation (Ashland, MA).

### LUV preparation

Large unilamellar vesicles (LUVs) were prepared using non-fluorescent and fluorescent phospholipids. Non-fluorescent lipids included DOPC (PC), DOPE (PE), and Soy-PI (PI), mixed at 65% PC, 27% PE, and 8% PI by weight in chloroform to mimic an LD monolayer, as previously done (15, 33). The fluorescent lipid, NR12S, was added at 0.1% w/w of the total phospholipids. Neutral lipids of triolein (TO), diolein (DO), or cholesteryl ester (CE) were added to the MLVs at a concentration of 0%, 1%, 5%, or 25% by weight of the total phospholipid content and below the concentration at which separate lipid droplets or droplet-embedded vesicles would be expected to form. The lipids were mixed with excess chloroform, dried under a nitrogen stream, kept under vacuum for at least 30 min, and resuspended in intracellular buffer (IB: 10 mM HEPES, 140 mM KCl, 6 mM NaCl, 1 mM MgCl₂, 2 mM EGTA, pH 7.4) to yield multilamellar vesicles (MLVs). The MLVs and neutral lipids were vortexed and extruded 21 times through a 100-nm pore-size membrane filter using a liposome extruder (Avanti Polar Lipids) to generate LUVs of varying neutral lipid content.

### GUV preparation

Giant unilamellar vesicles (GUVs) were prepared via electroformation, as described previously (33, 34). The phospholipid composition used in these preparations was 65:27:8 PC:PE:PI by weight. Neutral lipids at 0%, 2.5%, 5%, 10%, or 25% by weight of total phospholipids were mixed with phospholipids in chloroform. Lipid mixtures were dried on indium tin oxide (ITO) plates under vacuum for at least 30 minutes. The GUV growth chamber was then assembled using two ITO plates separated by a silicon spacer and filled with less than 1 mL of 307 mM sucrose in mQ-water. The osmolarity of the sucrose solution was adjusted to match that of the IB (307 mOsm). Electroformation was carried out using a 10 Hz AC voltage at 4 Vpp for 3 hours at 55°C. GUVs were carefully transferred to Eppendorf tubes using a truncated pipette tip to avoid bursting and stored at 4°C (33, 35).

### DEV preparation

Droplet-embedded vesicles (DEVs) were prepared by mixing a neutral lipid emulsion with the GUVs. The oil emulsions were prepared by adding 5 µL of TO or DO to 63 µL IB and 7 µL mQ-water, vortexing for 10 sec, bath-sonicating for 10 sec, and repeating 5-sec cycles of vortexing and sonication until a cloudy mixture was observed. DO was warmed to 37°C prior to use to ensure complete homogenization. 3.3 μL of the GUVs, 46.7 μL of IB, and 30 µL of the oil emulsion were gently combined in an Eppendorf tube and incubated on a rotator for 5 min at room temperature to allow for DEV formation from fusion between the GUVs and oil droplets, as described previously (35).

### Sample preparation for imaging

Glass-bottom well plates were cleaned with ethanol and water before being dried under a nitrogen stream. To enhance adhesion and stabilize the GUVs and DEVs on the glass surface, the coverslips were treated with 5% w/w BSA for 5 min and rinsed three times with IB. This treatment facilitated the settling of vesicles for observation with inverted microscopy. For fluorescent imaging of GUVs with varying neutral lipid compositions, GUVs were first diluted in IB at a 1:15 ratio. Then, 90 µL of IB was added to each pretreated well of a 24-well glass-bottom plate, followed by the addition of 10 µL of the diluted GUVs. For fluorescent imaging of DEVs with varying neutral lipid and phospholipid compositions, 100 µL of IB was added to each pretreated well of a 24-well glass-bottom plate, followed by 10 µL of the DEVs using a truncated pipette tip to prevent vesicle bursting. ABHD5-CFP was added to the wells at a final concentration of 100 nM and incubated at room temperature for 1 hour prior to imaging. For LUV imaging, 50 µL of LUVs at a concentration of 0.5 mg/mL were directly added to clean wells of a 96-well glass-bottom plate.

### Protein purification and preparation

Baculoviruses for protein expression were prepared using the Bac-to-Bac expression system (Invitrogen). pFastBac1 constructs with His tags were generated for Klarsicht, ANC-1, and Syne Homology–enhanced yellow fluorescent protein (KASH-YFP), mABHD5-mCherry, and mABHD5-CFP using standard molecular biology methods and confirmed by sequencing. Bacmid DNAs were generated by transformation of DH10Bac E. coli (Invitrogen) with FastBac plasmids following the manufacturer’s protocol. Initial baculovirus stocks were generated by transfection of Sf9 insect cells with bacmid DNA using Cellfectin II reagent (Invitrogen) according to the manufacturer’s protocol, then were amplified by infection of Sf9 cells with the initial baculoviral stocks. Amplified baculoviral stocks were used to infect High Five insect cells, which were collected 48 hours after infection when cells were 80% viable. High Five cells expressing proteins were lysed by sonication in IB buffer containing 20 µg/mL leupeptin and pepstatin A.

Supernatants were incubated with TALON Cobalt beads (Takara) for 2 hours at room temperature and eluted from the beads using 50 mM sodium phosphate buffer (pH 7.4) containing 150 mM imidazole. Imidazole was removed from proteins by successive rounds of concentration and redilution with IB buffer using Centricon centrifugal filter units of the appropriate molecular weight cutoff. Protein concentrations were determined by BCA (Pierce), and protein purity was assessed by Coomassie-stained PAGE gels.

Wild-type COS-7 cells were grown on uncoated microscopy coverslips in 6-well plates and transfected with ABHD5-mCherry and KASH-YFP using standard methods. Coverslips were mounted in a 35 mm metal imaging dish (Attofluor Cell Chamber, Invitrogen) for confocal microscopy prior to rinsing with phosphate-buffered saline and imaging in 10 mM HEPES Krebs-Ringer buffer at pH 7.4 (Sigma). Images were deconvolved within the Fusion software (Andor) with default settings.

### Confocal microscopy

Confocal fluorescence imaging was performed using an inverted microscope (DMi8, Leica) equipped with a spinning disk confocal unit (Dragonfly 200, Andor), a 63× oil-immersion objective, and an sCMOS camera (Zyla, Andor). Image acquisitions were carried out using Fusion software version 2.3.0.44.

### Fluorescence lifetime imaging microscopy

Fluorescence lifetime imaging microscopy (FLIM) was performed on both PicoQuant and Abberior confocal microscopes. Fluorescence lifetimes of NR12S within TO and DO LUVs were acquired using an inverted confocal microscope (IX73, Olympus) equipped with a 1.45 NA, 100× oil-immersion objective lens (UPLSAPO100XO, Olympus) and confocal, time-resolved, single-photon detection (MicroTime 200, PicoQuant). The confocal unit comprised a 485 nm wavelength pulsed diode laser (LDH-D-C-485, PicoQuant) at a repetition rate of 20 MHz, a 75 μm diameter aperture, and a single-photon avalanche diode (Photon Counting Module, Excelitas). Excitation and emission were separated via a dichroic mirror (ZT473/594rpc, Chroma) and a long-pass filter (488LP, Semrock) preceding detection. Time-correlated single-photon counting electronics (HydraHarp400, PicoQuant) in T3 mode recorded the single-photon emission time relative to the excitation laser pulse.

Fluorescence lifetimes of NR12S within CE LUVs were recorded using an inverted confocal microscope (Ti2, Nikon) equipped with a 1.45 NA, 100× oil-immersion objective lens (MRD71970, Nikon) and confocal, time-resolved, single-photon detection (STEDYCON, Abberior Instruments). Samples were excited in confocal mode using a 488 nm pulsed diode laser operating at a modulation frequency of 40 MHz at 10% power. Fluorescence emission was collected through three spectral channels (505–550 nm, 575–625 nm, 650–700 nm) using single-photon avalanche diodes, and the confocal pinhole was set to 50 µm (0.91 AU at 650 nm). Photon streams were recorded with the system’s Time-correlated single-photon counting electronics (MultiHarp 150, PicoQuant), allowing photon arrival times relative to the excitation pulse to be captured for fluorescence lifetime analysis. Images were acquired with a pixel dwell time of 10 µs, line frequency of 25 Hz, and single-line accumulation.

We collected 70 to 80 15-sec measurements and analyzed each measurement using a custom Python script. For each measurement, the phasor coordinates (*G*) and (*S*) were calculated directly from the photon arrival times using the fit-free phasor approach. Phasor analysis provides a model-free representation of fluorescence decay behavior and therefore does not require fitting the decay to a predefined single- or multiexponential model (36). The phasor coordinates were calculated according to

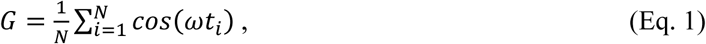

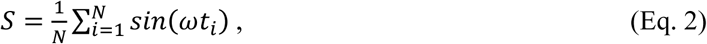

with the arrival time of the i^th^ photon (t_i_), the total number of photons (N), and the modulation angular frequency (*ω*). The fluorescence lifetime (*τ*) was calculated according to τ = S/(ωG).

### Molecular Dynamics Simulations

Coarse-grained models were constructed to investigate the effects of neutral lipids on lipid packing defects. We built phospholipid bilayers representing a conventional ER membrane and an LD-mimetic trilayer that consisted of two phospholipid monolayers with a neutral lipid core. The phospholipid bilayer and the phospholipid leaflets of the trilayer system consisted of POPC, DOPE, and SAPI at a mol ratio of 65:27:8. Neutral lipids were incorporated at concentrations defined relative to the total phospholipid content. For example, the 125% TO system contained 390 POPC, 162 DOPE, 48 SAPI, and 750 TO molecules. Initial coordinates for the lipid membranes were generated using CHARMM-GUI (37) in a bilayer configuration, and subsequent positioning of the neutral lipids between two phospholipid leaflets occurred spontaneously when the neutral lipid concentration was sufficiently high (15, 32).

All coarse-grained molecular dynamics simulations were performed using the MARTINI 3 force field (38) in the GROMACS 2018.4 simulation package (39). Systems were solvated with MARTINI water beads (38). Sodium and chloride ions were added to neutralize the system and achieve a physiological salt concentration of 0.15 M. Prior to equilibration, each system was minimized using two consecutive steps of 5,000 iterations, each using the steepest descent algorithm to remove unfavorable contacts and relax the initial configurations (40). The minimized structures were equilibrated through a multi-step process in which lipid positional restraints, starting with an initial restraint of 200 kJ/mol/nm² with integration time step of 2 fs, which was then reduced to 100, 50, 20, 10, and 10 kJ/mol/nm² and the time steps were increased to 5, 10, 15, 15, and 20 fs, respectively. The relative dielectric was set to 15, with long-range electrostatic and van der Waals interactions truncated at 1.1 nm using the reaction-field method. Temperature was maintained at 300 K using a velocity rescaling thermostat (41). Pressure was coupled semi-isotropically using Berendsen pressure coupling with a compressibility of 3 × 10⁻⁴ bar⁻¹, a reference pressure of 1 bar, and a coupling constant of 5 ps (42). For the production runs, the pressure coupling was changed to a Parrinello-Rahman barostat with a coupling constant of 12 ps, the same compressibility, and the same reference pressure (43). Production simulations were performed without positional constraints using a 20-fs time step. For each system, three independent replicas were simulated for 5 μs, and the last 2 μs of each trajectory were used for packing defect analysis.

The all-atom molecular dynamics (MD) simulations were reported previously and reanalyzed here (32, 44). Briefly, a coarse-grained MD simulation was used to construct the ABHD5-membrane complex based on the validated AlphaFold model of ABHD5 (45). This was followed by 50 ns of conventional all-atom MD for equilibration. Production sampling was then performed using Gaussian-accelerated MD (GaMD) for 1 µs with three independent replicas. All all-atom MD simulations were carried out using the Amber20 package with GPU acceleration (46). Simulations were conducted in the NPT ensemble at 303 K with 0.15 M KCl, a Langevin dynamics collision frequency of 5 ps⁻¹, and bonds involving hydrogen atoms constrained with the SHAKE algorithm. Long-range electrostatics were treated with the particle mesh Ewald (PME) method using a 12 Å cutoff, and the simulation time step was 2 fs.

### Lipid Packing Defect Analysis

Lipid packing defects were quantified using a custom, surface-based analysis workflow. For each coarse-grained molecular dynamics simulation trajectory frame, the membrane surface was reconstructed as a triangular mesh using the GL2 beads, which represent the glycerol groups of phospholipids. Unlike conventional grid-based approaches that rely on a flat reference plane (47, 48), this triangular mesh representation captures the local membrane geometry and preserves surface curvature. The mesh was generated at a resolution of 1 Å² per triangle. The upper and lower leaflets were identified and analyzed separately. For each mesh triangle, lipid coverage was evaluated based on the local surface normal and the surrounding phospholipid beads. A triangle was classified as a packing defect when less than 50% of its area was covered by phospholipid headgroup molecules. Adjacent defect triangles sharing common edges were grouped into connected defect clusters. Defect clusters arising from periodic boundary conditions were merged to eliminate duplicate counting. Small, isolated clusters corresponding to noise were removed, and narrow bridge-like connections between neighboring defects were pruned to separate physically distinct defect regions. The area of each defect cluster was calculated as the sum of the areas of all mesh triangles belonging to the cluster. The probability distribution of defect cluster areas (P(A)) was then determined from the ensemble of simulation frames. Packing defect constants (*π*) were obtained by fitting the defect area distribution to a single-exponential decay function (47, 49, 50), according to *P*(*A*) = *be*^-A/π^, where the pre-exponential factor (*b*) and the defect area (*A*) define the packing defect density. Larger values of *π* indicate a larger average defect size and increased membrane packing heterogeneity. A detailed analysis of this custom lipid packing defect analysis routine is freely available at (51). Simulation input files and trajectories have been deposited at (52).

## Results

To understand ABHD5 partitioning between different membrane environments, we visualized the localization of ABHD5 in COS-7 cells. ABHD5 was observed on LDs and the ER, binding to both phospholipid bilayers and monolayers (**Figs. 1A and S1**). To directly test the biophysical basis of this protein sorting, we utilized droplet-embedded vesicles (DEVs) (33, 35) as a reductionist model. DEVs provide a defined system with a phospholipid bilayer continuously connected to a phospholipid monolayer that surrounds an embedded neutral lipid core (**Fig. 1B**). DEVs allow for the simultaneous visualization of protein sorting between monolayer and bilayer surfaces. To complement these experiments, we integrated computational modeling (**Fig. 1C**) to resolve lipid behaviors and protein-membrane interactions within these distinct environments.

**Figure 1:**
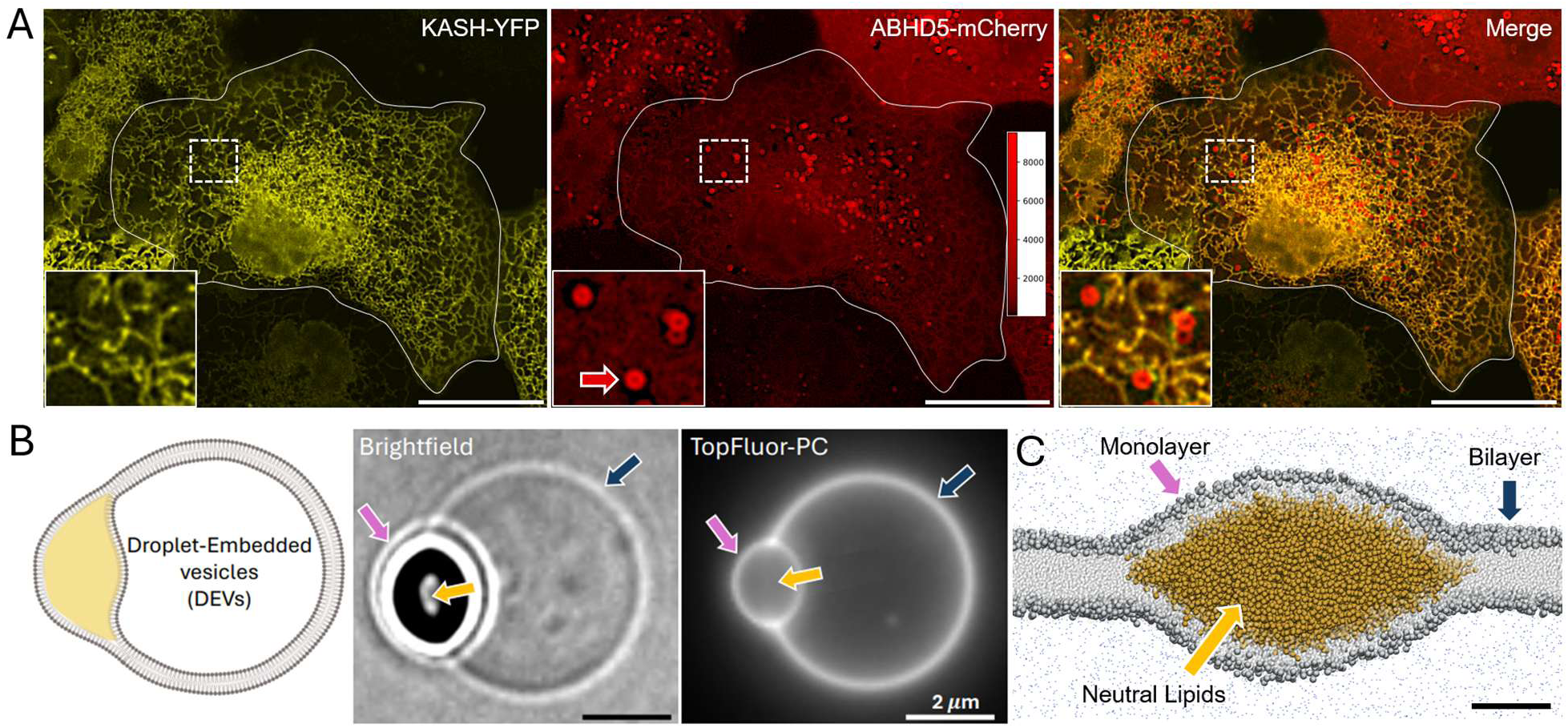
ABHD5 sorts between intracellular membranes and model systems used in this study. (**A**) Representative fluorescence images of live COS-7 cells, including the ER labeled with KASH-YFP (*left*), ABHD5-mCherry localizing to both LDs and the ER (*center*), and their merge (*right*). A single cell is outlined. The insets show an enlarged view of the regions indicated (*dashed boxes*), with an example round LD with concentrated ABHD5-mCherry (*red arrow*). An intensity line profile further demonstrates the binding of ABHD5 to the ER (Fig. S1). The scale bar represents 20 μm. (**B**) A schematic (*left*), a brightfield image (*center*), and a fluorescence image (*right*) of a DEV are shown. Model membrane DEVs allow simultaneous visualization of proteins associated with phospholipid bilayers (*blue arrows*), monolayers (*pink arrows*), and neutral lipids (*yellow arrows*) to quantify protein binding and sorting. (**C**) A coarse-grained molecular dynamics simulation with phospholipids and triolein was performed to show the partitioning of the neutral lipids between the phospholipid leaflets, as occurs in DEVs. The scale bar represents 5 nm.

### Neutral lipids affect the binding and sorting of ABHD5

We investigated the binding and sorting of ABHD5 to monolayers and bilayers of DEVs by varying the neutral lipid compositions, including 100% DO, 100% TO, and a 50:50 v/v mixture of DO and TO (**Fig. 2A**). Prior to ABHD5 addition, DEVs across all neutral lipid compositions were indistinguishable via brightfield. No observable differences in vesicle size, morphology, or stability were seen dependent upon the neutral lipid species incorporated. Further, no differences in DEV yield and emulsion stability were observed between neutral lipids. For all neutral lipids tested, ABHD5 bound more strongly to the DEV monolayer than the DEV bilayer; however, the ABHD5 binding strength and sorting depended on the specific neutral lipids present (**Fig. 2C**).

**Figure 2:**
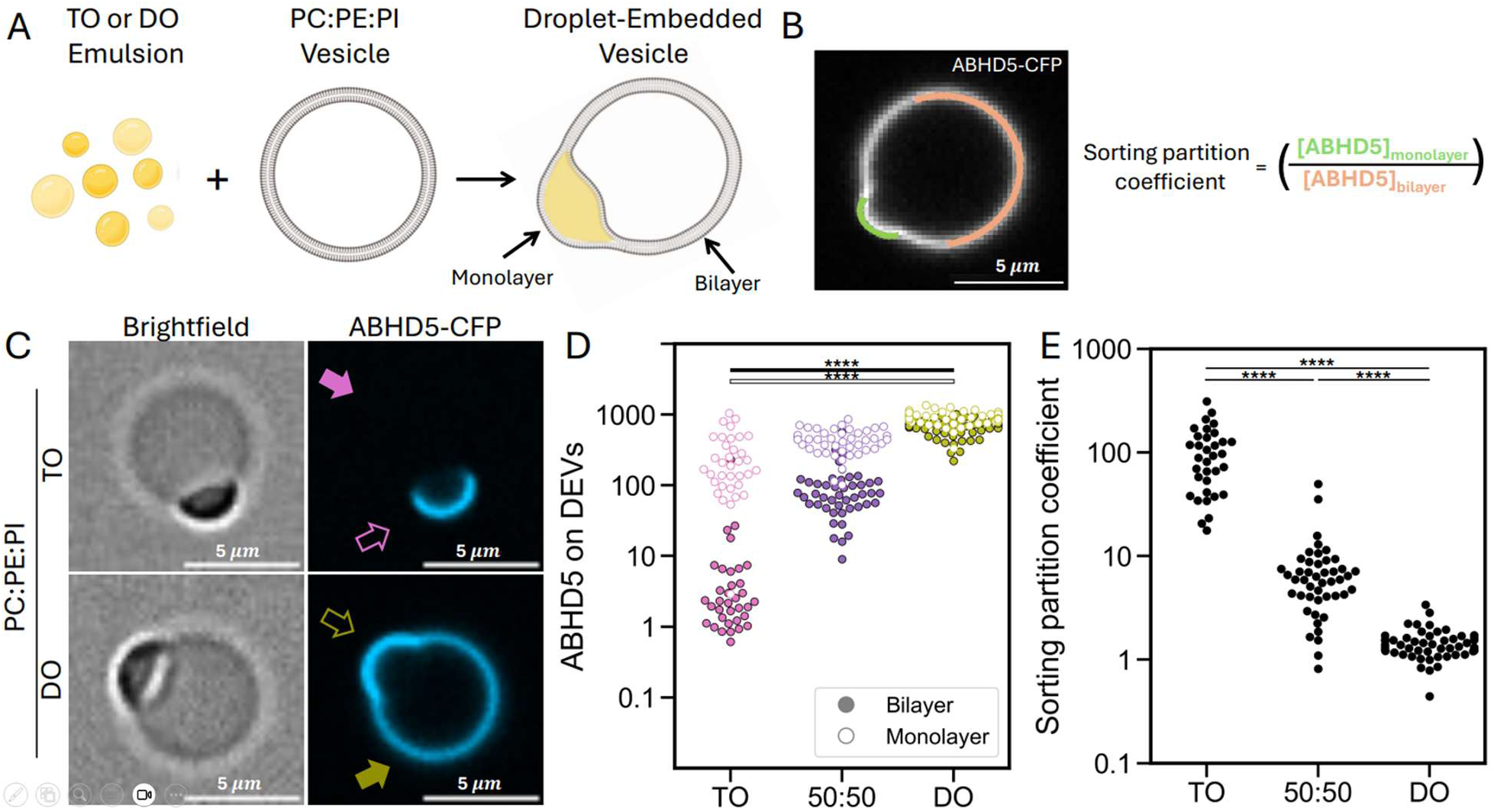
Neutral lipids affect the binding and sorting of ABHD5 in vitro. (**A**) Schematic showing DEV generation by mixing an emulsion containing diolein (DO), triolein (TO), or a 50:50 v/v mixture of DO and TO with GUVs composed of PC:PE:PI at a 65:27:8 weight ratio. (**B**) Representative fluorescent image of ABHD5-mCherry on a DO DEV highlighting the monolayer and the bilayer. The sorting partition coefficient was calculated as the ratio of ABHD5 intensity on the monolayer to the bilayer and displayed in 2E on a logarithmic scale. (**C**) Representative images of DEVs with TO (*top*) or DO (*bottom*) showing how the sorting varies based on the neutral lipids. (*D*) ABHD5 preferentially binds more to monolayers of TO DEVs; however, DEVs with DO show significantly greater ABHD5 binding to both the phospholipid bilayer and the monolayer. (**E**) Quantification of protein sorting between membranes shows a nearly 68-fold change in the partition coefficient with alterations in neutral lipid compositions. Filled and hollow significance bars represent bilayer and monolayer conditions, respectively. Statistical analysis was performed using Welch’s t-test; ****p < 0.0001.

For DEVs with 100% TO neutral lipids, we observed strong ABHD5 binding to the DEV monolayer and very weak binding to the DEV bilayer. As DO concentrations increased, ABHD5 binding increased in both the monolayers and bilayers (**Fig. 2D**). Furthermore, the relative sorting of ABHD5 between the monolayers and bilayers was influenced by the neutral lipid composition. Increasing the DO concentration resulted in more equal binding of ABHD5 to the monolayers and the bilayers, thereby demonstrating less sorting to the monolayer than observed on the TO DEVs.

To quantify, we measured the intensity of ABHD5 fluorescence on each DEV surface for each neutral lipid composition and calculated the sorting partition coefficient as the ratio of ABHD5 intensity on the monolayer to that on the bilayer (**Fig. 2B**). A lower value of the sorting partition coefficient indicates the protein was more evenly distributed between monolayer and bilayer; a higher value indicates the protein is more concentrated in the monolayer than in the bilayer. The TO DEVs yielded a sorting partition coefficient of 99 ± 12 (mean ± SEM), which was 68-fold higher than that of the DO DEVs at 1.45 ± 0.06 (mean ± SEM) (**Fig. 2E**).

### Computational analysis indicates that neutral lipids induce phospholipid packing defects

As a first attempt to understand the partitioning of neutral lipids in DEVs, we employed coarse-grained molecular dynamics simulations (**Fig. 3**). Using an experimentally consistent mixture of PC, PE, and PI phospholipids at a mol ratio of 65:27:8, we added varying concentrations of neutral lipids and computationally calculated the phospholipid packing defects. When TO and DO exceeded their solubility within the phospholipid bilayer, they separated into lipid droplets when the neutral lipid:phospholipid ratio exceeded 2 and 25%, respectively. Below these concentrations, the simulations revealed how neutral lipids affected phospholipid packing in a bilayer. The packing defects observed with the 1% TO system exceeded those with the 1% DO system, consistent with the larger TO molecular size and the greater disruption of phospholipids caused by one TO molecule versus one DO molecule. However, at greater DO concentrations, the DO remained soluble within the bilayer, and the DO-induced packing defects continued to increase beyond those ever created by TO in the bilayer (**Fig. 3F, G**). Greater DO packing defects were observed just prior to DO LD nucleation than the TO-generated defects in the bilayer prior to TO LD nucleation.

**Figure 3:**
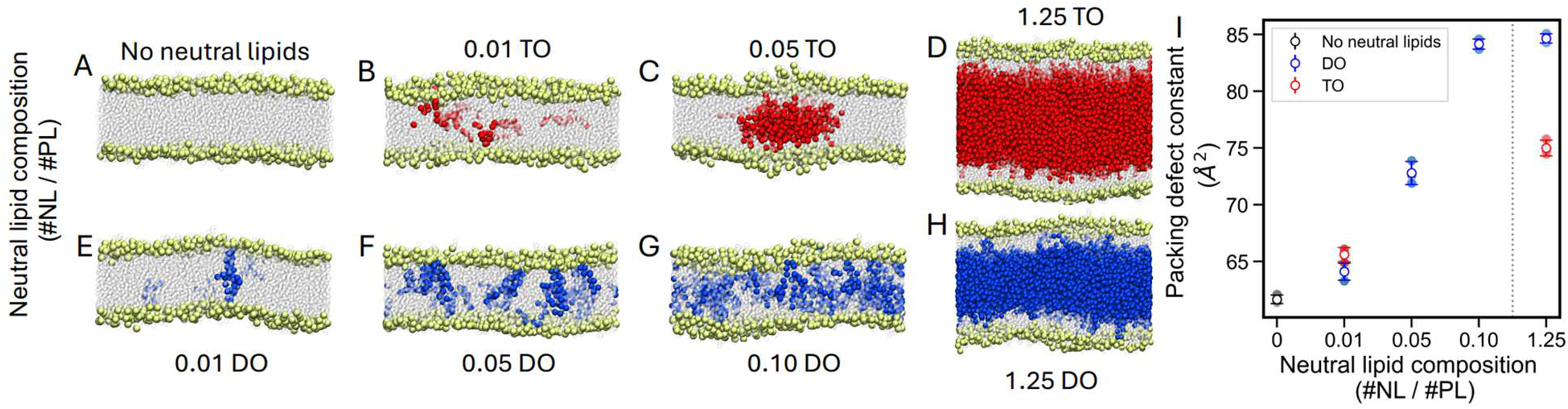
Coarse-grained molecular dynamics simulations reveal that neutral lipids induce phospholipid packing defects. (**A-H**) Computational simulation snapshots of phospholipid bilayers composed of PC:PE:PI (65:27:8 mol ratio) with varying concentrations of triolein (TO, *red*) and diolein (DO, *blue*). At 1%, neutral lipids were soluble in the phospholipid bilayer. DO remained dispersed in the bilayer at higher concentrations than TO. DO and TO exceeded their solubility limit and separated into lipid droplets at 2 and 25%, respectively. (**I**) The packing defect constant quantifies how DO and TO penetrate the phospholipid bilayer tails and generate packing defects, and more so for DO than TO. Filled circles represent the values from each simulation repeat, and hollow circles represent the mean ± SD. The gray dotted line visually separates the bilayer and monolayer regions.

When the concentration of neutral lipids exceeded their solubility within the bilayer, an LD nucleated and coexisted with a bilayer (**Fig. 3C**). As the neutral lipid concentration increased further, a trilayer system was generated with two phospholipid monolayers coating a neutral lipid core (**Fig. 3D, H**). Both the DO and TO penetrated the phospholipid monolayer tails and generated greater phospholipid packing defects than neutral-lipid-free phospholipid bilayers, with DO yielding 80% more packing defects than TO (**Fig 3I**). Interestingly, the TO monolayers had a similar density of packing defects as the near-saturating DO bilayers, and the TO DEV monolayers had similar ABHD5 binding as the DO DEV bilayers, which supports the conclusion that ABHD5 binding to phospholipid surfaces depends on the phospholipid packing defects.

### Neutral lipids experimentally increase bilayer defects

To experimentally examine the biophysical consequences of neutral lipid incorporation into phospholipid bilayers, we measured the fluorescence lifetime of NR12S, a reporter of phospholipid packing (53, 54). A reduced NR12S fluorescence lifetime is generally consistent with lower membrane organization and reduced lipid packing, likely because the Nile Red chromophore experiences a more hydrated, polar, and dynamically relaxed interfacial environment, which increases nonradiative decay and shortens the observed excited-state lifetime (53, 54). In our system, the lipids all contained oleoyl acyl chains, so changes in NR12S lifetime would not be expected to arise primarily from differences in acyl-chain saturation or length. However, TO and DO are expected to perturb interfacial phospholipid packing due to their large hydrophobic volume and small or absent polar headgroups (i.e., their large effective packing parameters) and increase the NR12S fluorescence lifetime accordingly.

To measure the impact of neutral lipids on bilayer lipid packing, we incorporated neutral lipids into LUV bilayers at concentrations below their solubility limit, which are unlikely to form DEVs, and measured the fluorescence lifetime of NR12S (**Fig. 4A**). LUVs with DO and CE were made at 1%, 5%, 10%, and 25% by weight of the total phospholipid content. Because the TO forms DEVs at lower concentrations, LUVs with TO were only made at a 1% w/w TO-to-phospholipids ratio. Whereas DEVs provided an equilibrated partitioning of neutral lipids between the bilayer and the embedded LD, these LUV measurements enabled titration of neutral lipids in the bilayer.

**Figure 4:**
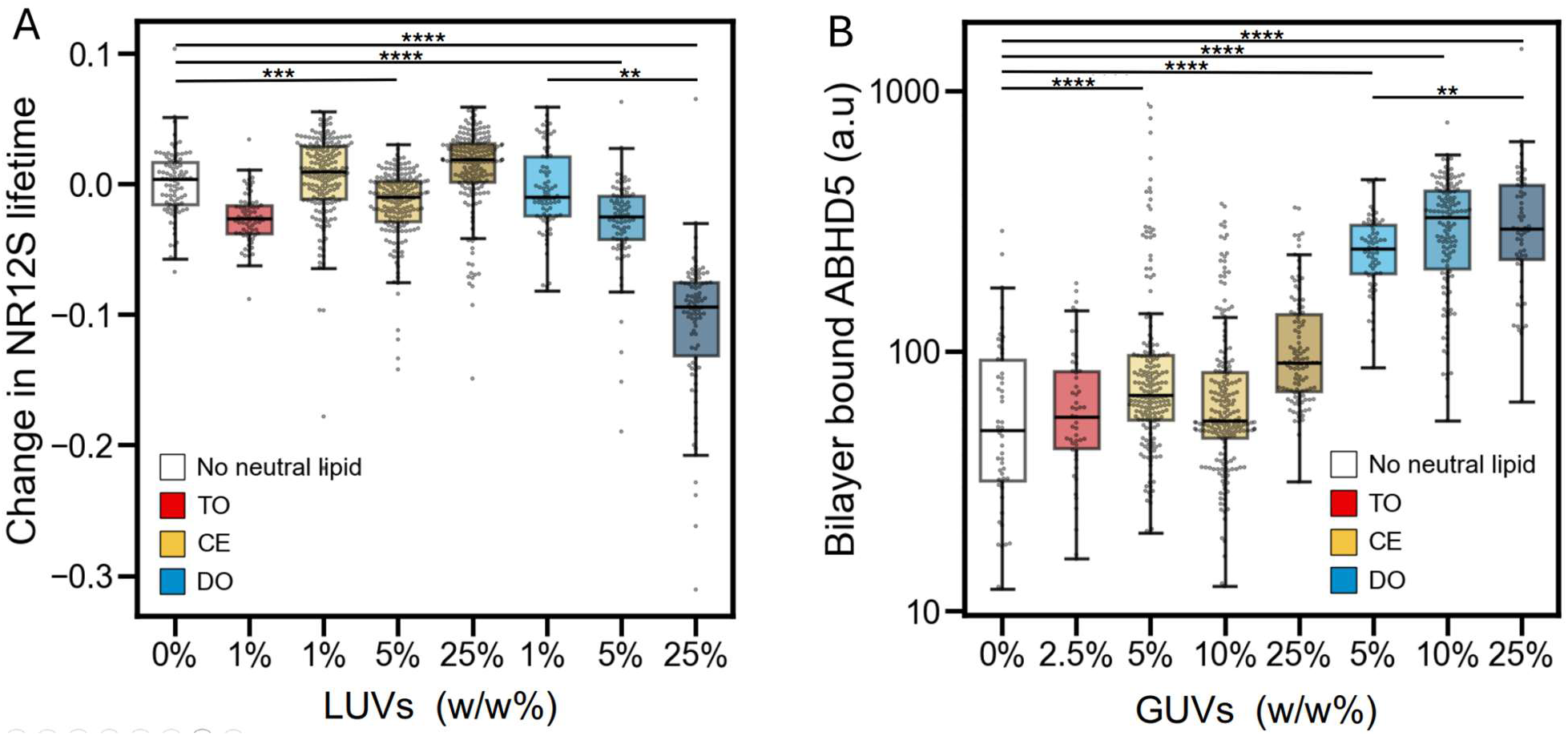
Neutral lipid increases bilayer disorder as reported by fluorescence lifetime microscopy and the effect of neutral lipid composition on ABHD5 binding to GUVs. (**A**) The change in NR12S lifetime was measured in LUVs containing triolein (TO), cholesteryl ester (CE), diolein (DO), or no neutral lipids relative to no-neutral-lipid LUVs. Individual circles represent one 15-sec measurement segment. Each experiment was performed at least three times, with measurements collected across three individual wells per experiment. Increasing DO generated the largest reduction in NR12S lifetime, whereas CE and TO produced comparatively smaller changes. (**B**) ABHD5 binding was quantified on phospholipid GUVs containing varying neutral lipids. Individual circles represent measurements from single GUVs. Each experiment was performed at least three times, with measurements collected across three individual wells per experiment. ABHD5 binding is displayed on a logarithmic scale. CE-containing GUVs (5% and 25%) and all DO-containing GUVs exhibited significantly greater ABHD5 binding than GUVs without neutral lipids, and ABHD5 binding was significantly higher on 25% DO GUVs than on 5% DO GUVs. Statistical significance was assessed using Welch’s t-test. Statistical significance is indicated as *p* < 0.01 (**), *p* < 0.001 (***), and *p <* 0.0001 (****).

The most dramatic effects of phospholipid packing were observed with DO incorporation. The fluorescence lifetime of NR12S decreased as DO concentration increased, indicating that the bilayer had more packing defects upon DO incorporation. However, consistent with the simulations, the addition of 1% w/w TO increased the packing defects more than 1% w/w DO, and no significant change was observed for CE incorporation.

### Neutral lipids increase ABHD5 binding to phospholipid bilayers

To further see the ramifications of ABHD5 binding to varying the neutral lipid content of bilayers that were primarily composed of phospholipids, we quantified ABHD5 binding to GUVs containing different concentrations of TO, CE, or DO (**Fig. 4B**). Incorporation of 2.5% TO did not significantly alter ABHD5 binding compared with GUVs with no neutral lipids. GUVs containing 5% or 25% CE exhibited significantly greater ABHD5 binding than neutral-lipid-containing GUVs, respectively, whereas the difference between no neutral lipid and 10% CE was not significant. ABHD5 binding also did not differ significantly between 5% and 25% CE. Thus, CE incorporation enhanced ABHD5 binding at some concentrations, but the data did not reveal a consistent concentration-dependent relationship. In contrast, GUVs containing 5%, 10%, or 25% DO displayed significantly greater ABHD5 binding than GUVs with no neutral-lipid content. Furthermore, binding was significantly greater for 25% DO than for 5% DO, supporting increased ABHD5 association with increasing DO content. Together, these results indicate that ABHD5 membrane association is sensitive to bilayer composition. TO had no detectable effect, whereas CE enhanced ABHD5 binding at 5% and 25% without showing a clear concentration-dependent pattern. DO produced the strongest and most consistent enhancement of ABHD5 binding, and the significant difference between 5% and 25% DO supports a concentration-dependent effect of DO enrichment.

### Phospholipid-dependent ABHD5 binding to monolayers and bilayers

To assess the effects of phospholipids on ABHD5 binding and sorting, we created DEVs with four phospholipid compositions: PC alone, PC:PE (73:27), PC:PE:PI (65:27:8), and PC:PE:PS (65:27:8) by weight. The fluorescence intensity of ABHD5-CFP was measured on DEV monolayers and bilayers to assess ABHD5 binding and sorting as a function of the combined effects of phospholipids and neutral lipids in forming the lipid surfaces.

When the neutral lipid composition was TO only (**Fig. 5A**), ABHD5 binding was lowest in DEVs composed of PC alone. Monolayer-associated fluorescence increased significantly following the addition of PE and remained elevated with the incorporation of PI or PS, whereas no significant differences were observed among the PC:PE, PC:PE:PI, and PC:PE:PS compositions. In contrast, bilayer-associated ABHD5 fluorescence remained relatively constant across the different phospholipid compositions, with the only significant difference detected between the PC:PE:PI and PC:PE:PS conditions. Across all phospholipid compositions, ABHD5 consistently associated more strongly with the monolayer than the bilayer. However, the sorting partition coefficient did not differ significantly among the four phospholipid compositions for TO DEVs (**Fig. 5B**), indicating that the relative enrichment of ABHD5 on the monolayer compared with the bilayer was maintained regardless of phospholipid composition.

**Figure 5:**
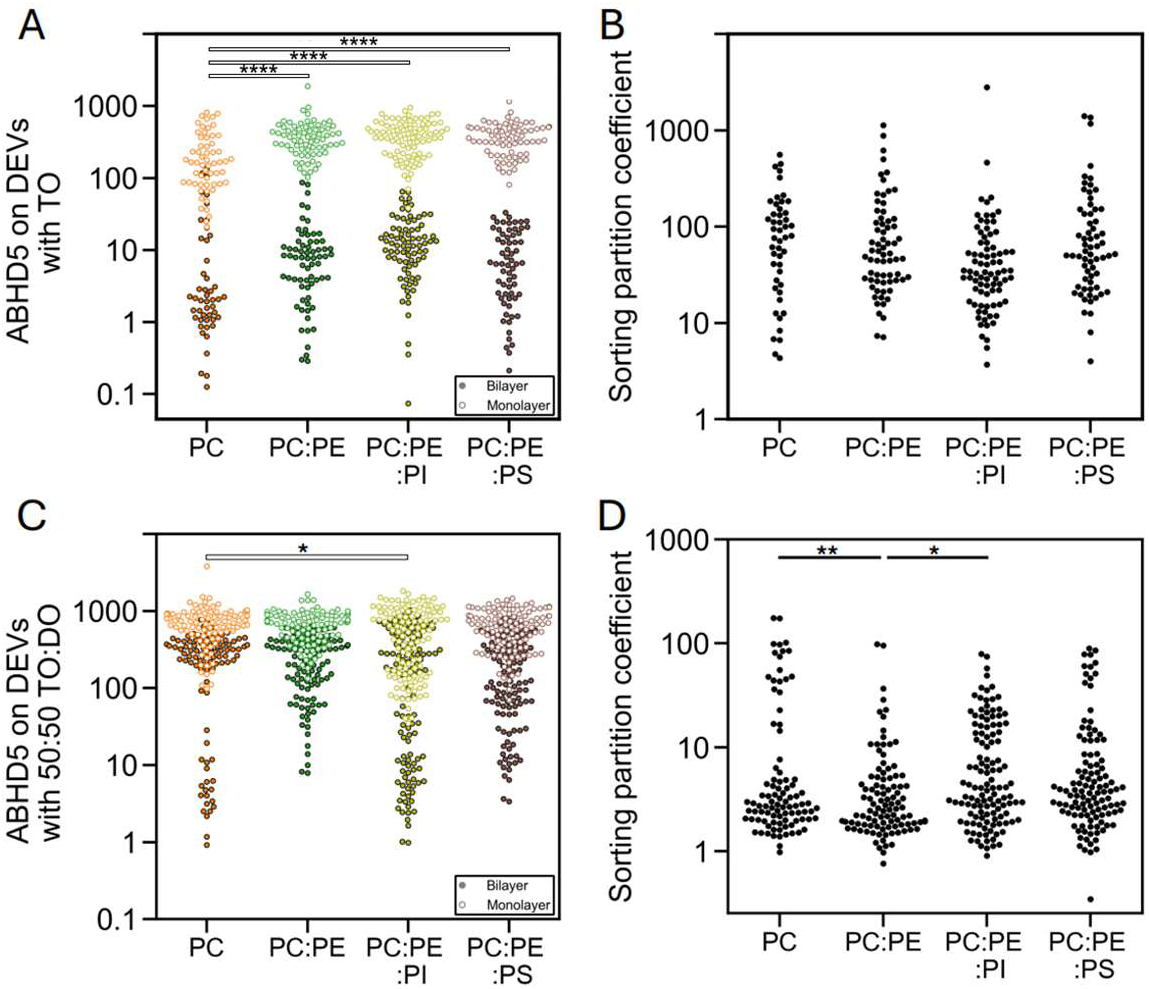
Phospholipid-dependent ABHD5 binding and sorting. ABHD5 binding and sorting on DEVs with (**A, B**) TO or (**C, D**) 50:50 v/v TO:DO were measured across varying phospholipid compositions, including PC alone, PC:PE (73:27 w/w), PC:PE:PI (65:27:8 w/w), or PC:PE:PS (65:27:8 w/w). ABHD5 preferentially bound more to the monolayer across all phospholipid and neutral lipid types. For the TO DEVs, no significant differences were detected in bilayer binding between the phospholipid compositions. However, the TO DEVs of only PC demonstrated less monolayer binding than all other phospholipid compositions. For the TO:DO DEVs, ABHD5 fluorescence remained high across all phospholipid compositions; monolayer- and bilayer-associated fluorescence was consistently higher for TO:DO DEVs than for TO DEVs. Individual circles represent single DEV measurements. Data were collected from at least five different wells for each of at least five repeated experiments. Statistical significance was assessed using Welch’s t-test and indicated as *p* < 0.05 (*), *p* < 0.01 (**), *p* < 0.001 (***), *p <* 0.0001 (****).

In DEVs with TO and DO at 50:50 v/v (**Fig. 5C**), ABHD5 binding remained high across all phospholipid compositions in both monolayers and bilayers. Relative to PC, a significant increase in fluorescence was observed only for the monolayers of PC:PE:PI composition, whereas the remaining comparisons were not statistically significant. Again, the monolayer-associated fluorescence remained greater than bilayer-associated fluorescence under all conditions. However, compared with the TO-only-DEVs, bilayer-associated ABHD5 fluorescence was consistently higher across all phospholipid compositions, resulting in a smaller difference between bilayer and monolayer fluorescence.

Although overall ABHD5 recruitment differed among phospholipid compositions, changes in the sorting partition coefficient in the mixed TO and DO DEVs were modest (**Fig. 5D**). Significant differences were detected only between PC and PC:PE and between PC:PE and PC:PE:PI, while all other comparisons were not statistically significant. We also observed that the presence of DO was associated with higher ABHD5 fluorescence on both the bilayer and monolayer across all lipid compositions compared with the TO-only condition (**Fig. 5A, B**), suggesting greater packing defects when DO was included in the DEVs.

## Discussion

### Lipid surface architecture regulates ABHD5 recruitment

Beyond the binding of ABHD5 to PLINs on LDs, recent data have shown that ABHD5 directly targets the ER and LDs depending on the cellular state, suggesting that the intracellular targeting of ABHD5 is influenced by the physical properties of lipid surfaces. This work directly addresses how lipid properties influence ABHD5 binding by using complementary experimental model systems and molecular simulations. We found that ABHD5 preferentially associates with lipid surfaces that contain packing defects. These findings indicate that changes in lipid composition and phospholipid organization may create favorable environments for ABHD5 recruitment.

Our data show how the physical differences between LD monolayers and ER bilayers contribute to ABHD5 recruitment. These lipid surfaces differ fundamentally in their architecture, including packing density, curvature, interfacial organization, and surface tension (13, 15, 16). LD monolayers contain a higher density of lipid packing defects than phospholipid bilayers, providing more favorable sites for protein association (14). Consistent with these differences, ABHD5 preferentially associated with LD monolayers rather than phospholipid bilayers, with a stronger dependence on the neutral lipid composition than the phospholipid composition among those examined (**Fig. 5**). This observation suggests that the intrinsic physical properties of LD monolayers provide a more favorable environment for ABHD5 recruitment than those of phospholipid bilayers. In addition to the intrinsic differences between LD monolayers and ER bilayers, changes in the cellular neutral lipid and phospholipid compositions would provide a mechanism for regulated intracellular trafficking of ABHD5.

### Mechanisms of lipid surface recognition

The interaction of ABHD5 with lipid surfaces can be explained by the two structurally distinct membrane-binding regions that contain hydrophobic residues and amphipathic helices of varying stability (32, 55). The largely unstructured ABHD5 N-terminal region (residues 1-43) serves as the initial lipid-surface sensor. It contains aromatic residues, including W21, W25, and W29, that are required for LD association (55). Following this initial docking step, the insertion segment (residues 180-230) engages the lipid surface, stabilizing the ABHD5-lipid interaction. This region contains the lid helix (residues 198-207) and several bulky hydrophobic residues, including W199, F210, and F222, that form a hydrophobic patch important for lipid surface association (32). Together, these features support a two-step mechanism in which ABHD5 first docks at the lipid surface and then stabilizes its association through hydrophobic insertion. This structural model provides a plausible explanation for the increased ABHD5 recruitment observed in defect-rich lipid surfaces in our study.

The lipid surface interaction mechanism of ABHD5 shares characteristic binding motifs with other lipid surface-binding proteins, including bulky, hydrophobic residues and amphipathic helices. Amphipathic helices typically mediate shallow interfacial interactions and are highly sensitive to lipid packing, curvature, and packing defects (11, 56). PLIN3 is a well-characterized LD-associated protein that binds LD monolayers through amphipathic helices (4, 57–59), and iPLA₂ uses an amphipathic helix to recognize membranes enriched in packing defects (60). Amphipathic lipid packing sensor motifs preferentially associate with defect-rich lipid surfaces (26, 48). Our findings suggest that ABHD5 follows the same general biophysical principle, as its recruitment increases at lipid surfaces with greater packing defects. ABHD5 exhibits a similar preference for defect-rich lipid surfaces that may allow ABHD5 to respond differently to changes in lipid surface organization via specific hydrophobic and amphiphilic motifs described below.

A deeper assessment of the helical motifs responsible for ABHD5-lipid association was performed by analyzing the secondary structure of ABHD5 using the compute_dssp function in the MDTraj package for the three 1 *μ*s all-atom molecular dynamics simulation replicas of LD- and ER-bound ABHD5, which were first reported and analyzed previously (32, 44). Regions that remained within an average distance of 0.5 nm from the phospholipid surface while adopting an α-helical conformation for at least 66% of the simulation included residues 35-43, 197-203, 213-216, 226-229, and 231-235 in the LD simulations, and residues 34-43, 185-187, 199-205, 212-215, and 232-235 in the ER simulations (**Figs. S3C, D and S4C, D**). Sequence analysis revealed that these membrane-associated motifs contained mixed hydrophobic, polar, and charged amino acids (**Fig. S5**), rather than extended hydrophobic stretches, suggesting chemically heterogeneous membrane-interacting regions.

LD monolayers contain more abundant lipid packing defects than bilayers, which have been shown to influence the membrane association of peripheral proteins and amphipathic membrane-binding motifs. Compared with the ER-bound ABHD5, the LD-bound ABHD5 exhibited a greater number of persistent membrane-associated helical motifs (**Figs. S3 and S4**), reflecting how the distinct physicochemical properties of lipid droplet monolayers influence the structure of bound proteins. The LD surface provides a more favorable environment for transient membrane interactions that are associated with localized helix formation in ABHD5.

The phospholipid-binding helices of ABHD5 are more hydrophobic than classical amphipathic helices, with their hydrophobic residues more broadly distributed rather than confined to one face of the helix (**Fig. S5**). This organization may allow deeper insertion into disordered lipid surfaces and a greater sensitivity to phospholipid packing defects that enhance its LD-specific targeting. Future studies directly quantifying lipid packing defects and residue-specific membrane interactions will be required to test this possibility.

### Neutral lipids modify lipid surface properties and promote ABHD5 recruitment

We further found that neutral lipids enhanced ABHD5 recruitment to both LD monolayers and phospholipid bilayers across multiple experimental model systems and molecular simulations, including before and after the neutral lipids condense into distinct LDs. Our simulations suggest that neutral lipids insert between phospholipid acyl chains, disrupting phospholipid packing and increasing lipid surface disorder (**Fig. 3**). This interpretation is supported by the reduced NR12S lifetime measured in LUVs, which is consistent with decreased lipid order and an increased prevalence of packing defects (**Fig. 4**). The increase in ABHD5 recruitment closely follows these changes in lipid surface organization, suggesting that neutral lipids promote protein association by increasing the availability of packing defects. Similar behavior has been reported for other lipid surface-binding proteins. For example, PLIN3 and amphipathic lipid packing sensor motifs exhibit enhanced association with diolein-containing bilayers through amphipathic helices (50, 57–59).

Together, these observations suggest that neutral lipids, and especially diolein, disrupt phospholipid packing through their conical geometry (61), increasing the availability of lipid packing defects and creating a more favorable environment for ABHD5 recruitment. In contrast to diolein, triolein influences lipid surface properties through a different mechanism. Triolein accumulation promotes phase separation into a neutral lipid lens, contributing to bilayer asymmetry and curvature stress (62). These changes alter the physical properties of lipid surfaces and are also expected to increase the availability of packing voids. Thus, although diolein and triolein remodel lipid surfaces with varying solubility in the bilayer, both generate defect-rich environments that may favor the recruitment of lipid surface-binding proteins, including ABHD5.

### Biological implications and future directions

Our findings support a model in which lipid surface properties contribute to ABHD5 localization by regulating the availability of lipid packing defects. Diacylglycerol production and accumulation in the ER should increase its packing defects and ABHD5 binding. To complement previous studies showing protein-mediated recruitment through perilipins (5, 6), our results suggest that ABHD5 localization may be guided by its direct lipid surface regulate recognition. These mechanisms may act cooperatively to regulate ABHD5 localization under different physiological conditions. While our study identifies lipid packing defects as an important determinant of ABHD5 recruitment, it does not establish how these mechanisms are integrated with cellular signaling pathways or dynamic changes in lipid droplet composition in vivo. Future studies will determine how lipid surface remodeling and protein-mediated interactions cooperate to regulate ABHD5 localization and function during lipid droplet formation, growth, and lipolysis.

## Data availability

Data generated in this study are provided in the Figures and the Supplementary Information. The all-atom molecular dynamics simulations presented and analyzed in the Supplementary Information were previously reported and are available at Zenodo.org (https://doi.org/10.5281/zenodo.15122721). A detailed description of the custom lipid packing defect analysis routine used in this study is freely available at Zenodo.org (https://doi.org/10.5281/zenodo.21907725). Simulation input files and trajectories are available at Zenodo.org (https://doi.org/10.5281/zenodo.21909038).

## Acknowledgements

We thank the Wayne State University High Performance Computing Center for providing resources for coarse-grained molecular dynamics simulations. The views expressed in this article are those of the authors and do not reflect the official policy or position of the U.S. Naval Academy, Department of the Navy, the Department of War, or the U.S. Government. Graphical abstracts were created with BioRender.com.

## Author contributions

Conceptualization and methodology: S.P. and C.V.K., project design: S.P. and C.V.K., investigation: A.N., A.K., Y.M.H., C.V.K. performed the simulations, Resources: M.A.S. and J.G.G., GUV and DEV data acquisition and imaging performed: S.P. and K.A., FLIM data acquisition and analysis performed: K.T., J.S., and S.P., manuscript writing: S.P., K. T., C.V.K.

---

**Figure S1:**
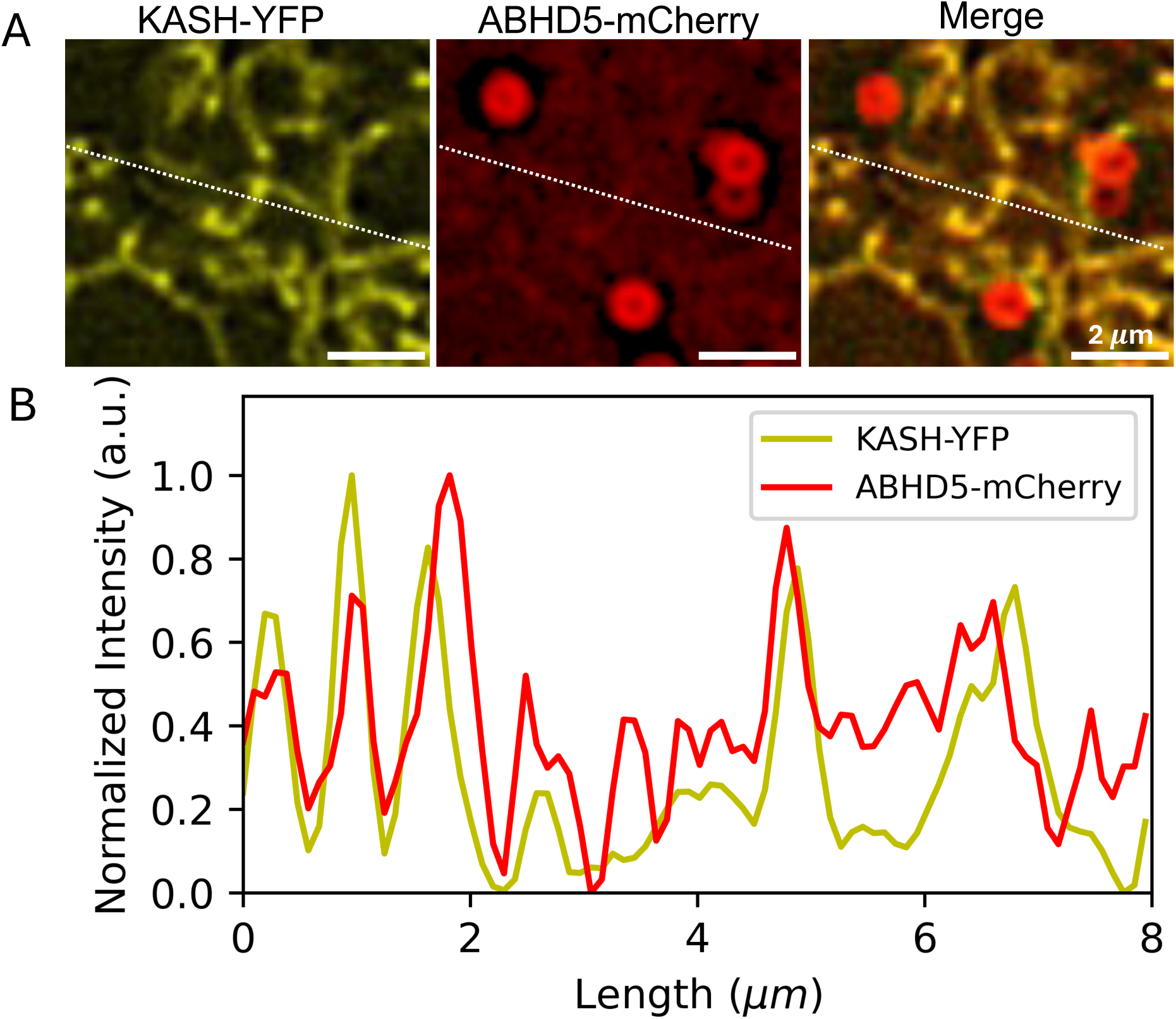
ABHD5 localizes to LDs and ER. (A) Enlarged view of the region indicated in the Figure 1A (right). Fluorescent image showing KASH-YFP (left), ABHD5-mCherry (center), and the merged channels (right). KASH-YFP labels the ER network, while ABHD5-mCherry is enriched on LDs and shows signal associated with the ER. The white dashed line indicates the region used for line-scan analysis. (B) Normalized fluorescence intensity profiles of KASH-YFP and ABHD5-mCherry along the indicated line scan. Overlapping peaks in KASH-YFP and ABHD5-mCherry fluorescence indicate spatial association of ABHD5 with the ER.

**Figure S2:**
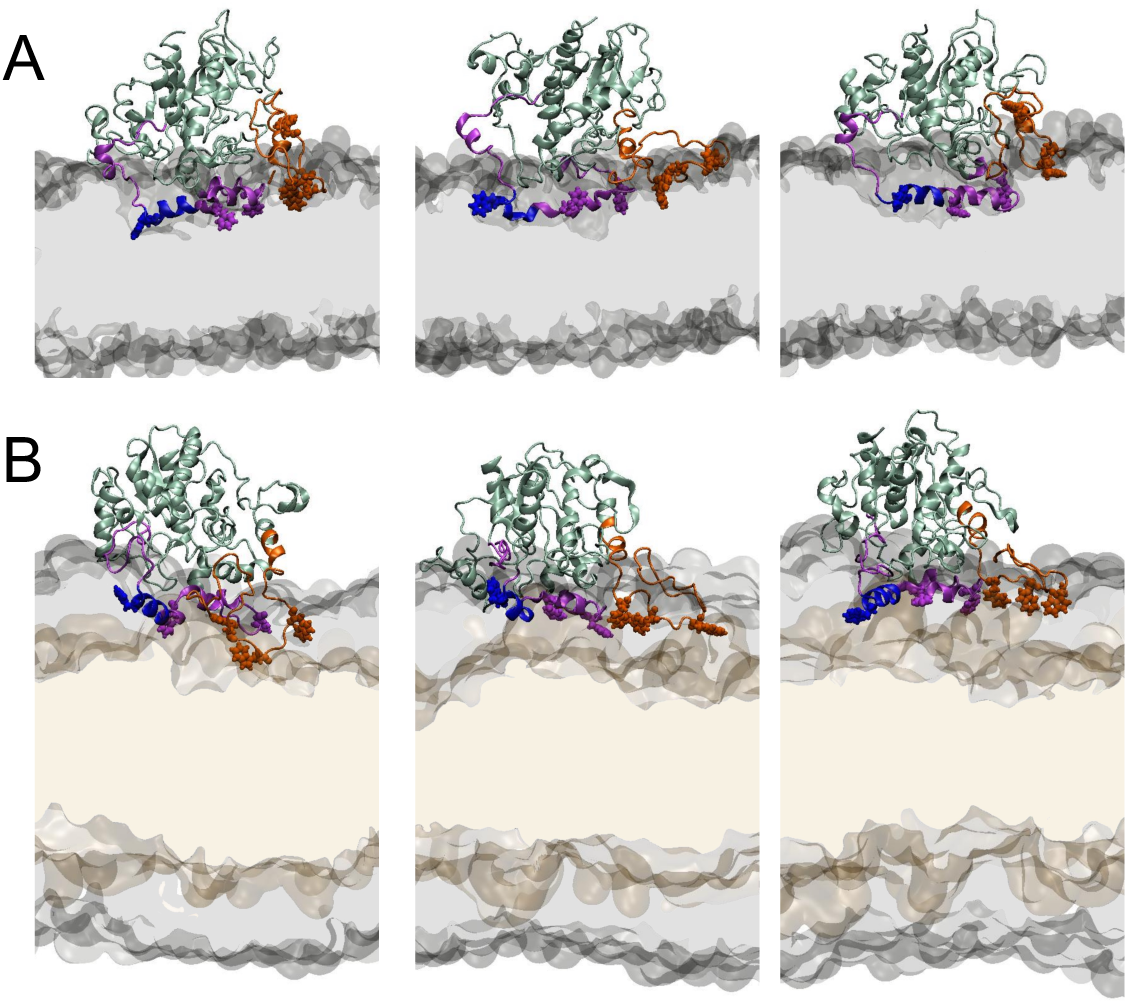
All-atom molecular dynamics snapshots of ABHD5 binding to (A) a phospholipid bilayer or (B) phospholipid monolayers surrounding a layer of triolein. The PC, PE, and PI phospholipids (*gray*) and neutral lipids (*brown*) attract the ABHD5 N-terminal region (aa1-43, *orange*) and the insertion domain (aa180-230, *purple*), which includes the lid (aa192-207, *blue*). W21, W25, W29, S199, F210, and F222 are bulky hydrophobic residues instrumental in lipid binding (*space-filling*). One view from each of the three replicas of each simulation is shown.

**Figure S3:**
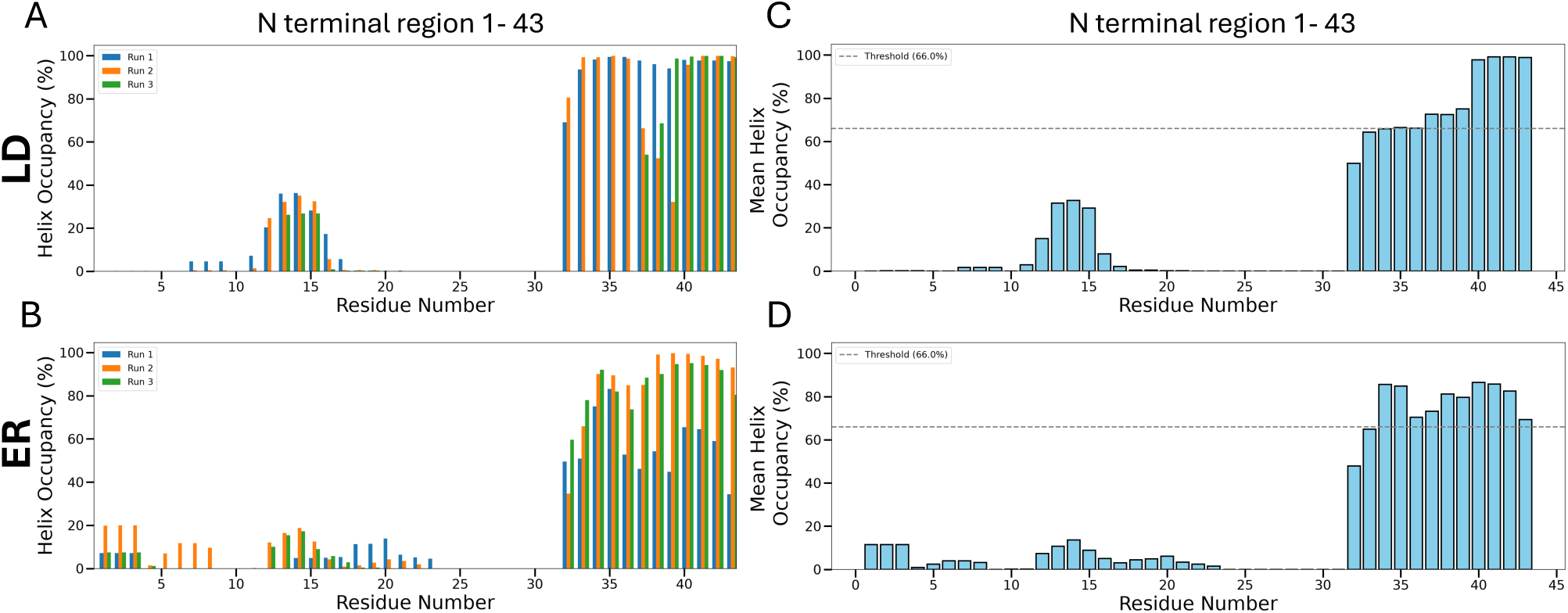
Helix occupancy was calculated for residues from the three repeated all-atom molecular dynamics simulations in the membrane binding regions 1-43.

**Figure S4.**
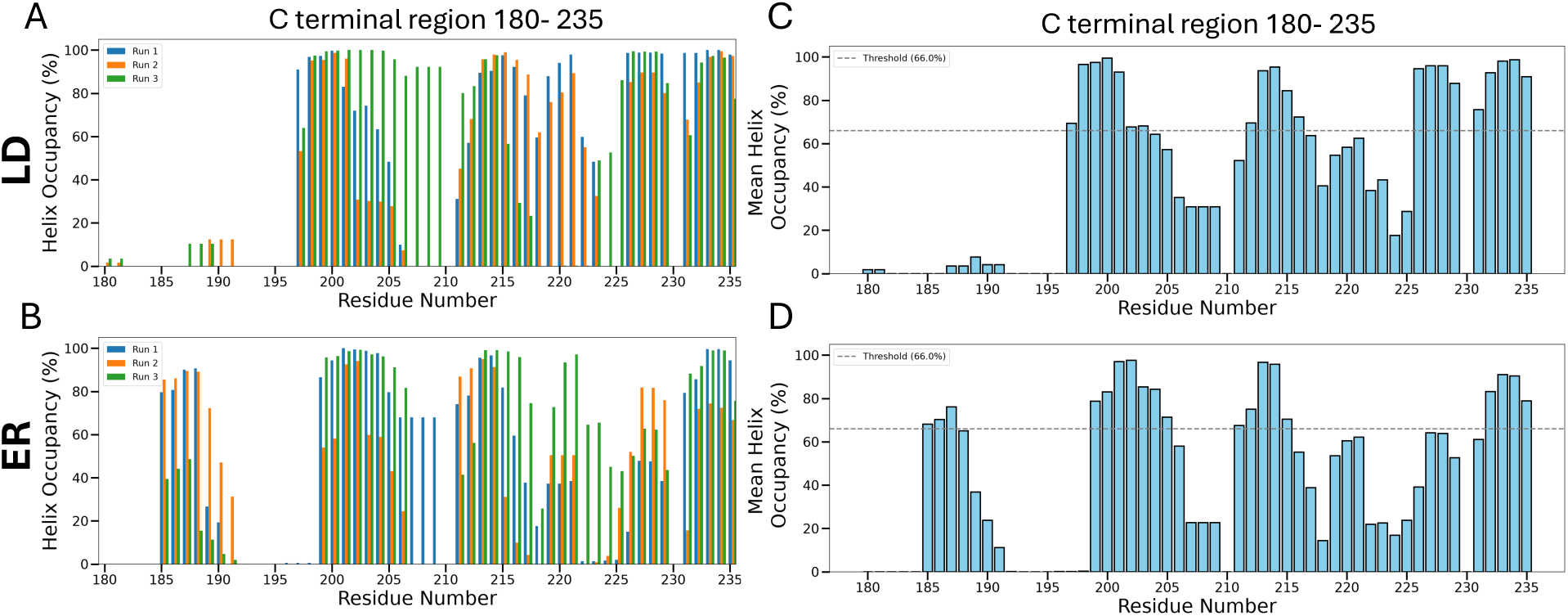
Helix occupancy was calculated for residues from the three repeated all-atom molecular dynamics in the membrane binding regions 180-235.

**Figure S5:**
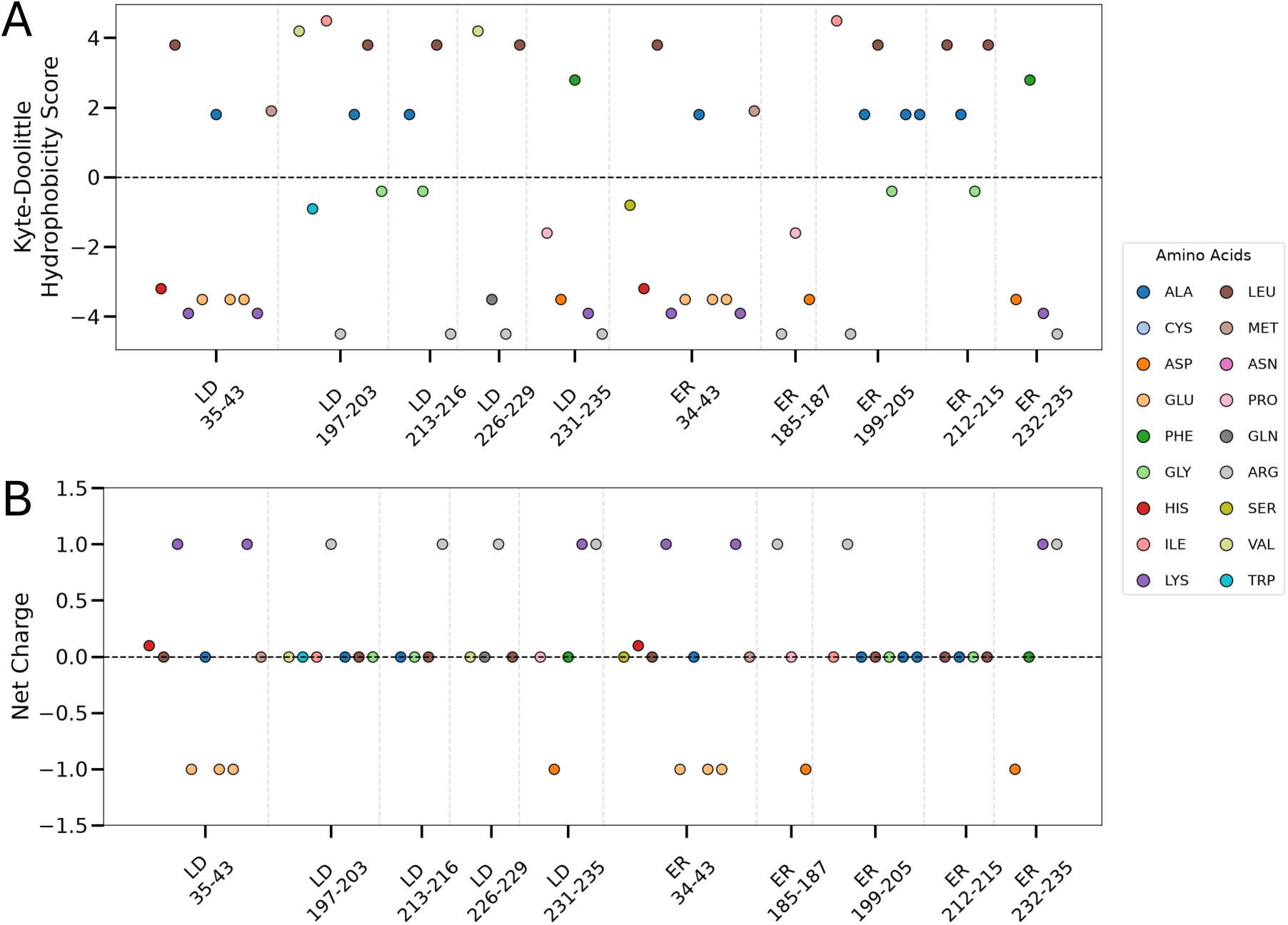
Physicochemical properties of persistent helical motifs support lipid-dependent membrane binding

**Figure S6:**
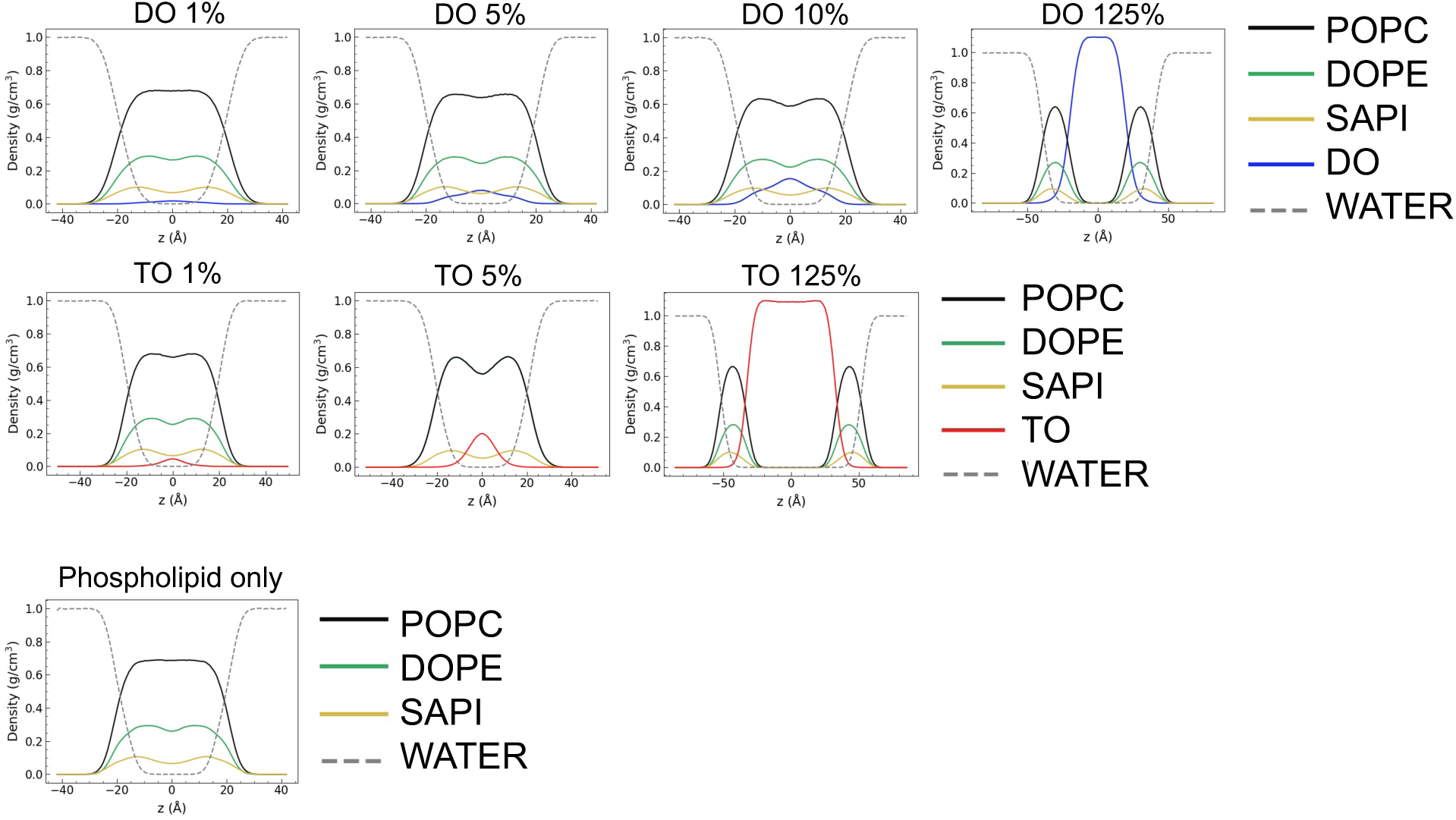
Z density profile for phospholipid and neutral lipids at varying concentrations.

**Table S1.** Pairwise comparison *p*-values for monolayer, bilayer, and sorting values across neutral lipid types (**Fig. 2**).

| **BILAYER vs BILAYER comparisons:** |  |  |
| --- | --- | --- |
| Pairs | *P* values | Summary |
| DO vs 50:50 | 2.236e-31 | **** |
| 50:50 vs TO | 3.240e-07 | **** |
| DO vs TO | 4.146e-32 | **** |
| **MONOLAYER vs MONOLAYER comparisons:** |  |  |
| Pairs | *P* values | Summary |
| DO vs 50:50 | 1.617e-23 | **** |
| 50:50 vs TO | 0.0019 | ** |
| DO vs TO | 4.007e-18 | **** |
| **SORTING PARTITION COEFFICIENT comparisons:** |  |  |
| Pairs | *P* values | Summary |
| DO vs 50:50 | 7.663e-06 | **** |
| 50:50 vs TO | 4.441e-09 | **** |
| DO vs TO | 1.041e-09 | **** |

**Table S2.** Pairwise comparison *p*-values for LUVs and GUVs with varying neutral lipid composition and concentrations (**Fig. 4**)

| **LUV Pairs** | ***P* values** | **Summary** |
| --- | --- | --- |
| 0% NL vs 1% TO | 4.641e-12 | **** |
| 0% NL vs 1% CE | 0.238 | n.s. |
| 0% NL vs 5% CE | 6.5411e-06 | **** |
| 0% NL vs 25% CE | 0.0013 | ** |
| 0% NL vs 1% DO | 0.265 | n.s. |
| 0% NL vs 5% DO | 5.385e-09 | **** |
| 0% NL vs 25% DO | 3.682e-35 | **** |
| 5% DO vs 25% DO | 7.405e-22 | **** |
| 5% CE vs 25% CE | 8.719e-17 | **** |

| **GUV Pairs** | ***P* values** | **Summary** |
| --- | --- | --- |
| 0% NL vs 2.5% TO | 0.996 | n.s. |
| 0% NL vs 5% CE | 0.0002457 | *** |
| 0% NL vs 10% CE | 0.187 | n.s. |
| 0% NL vs 25% CE | 4.827e-06 | **** |
| 0% NL vs 5% DO | 6.403e-28 | **** |
| 0% NL vs 10% DO | 2.403e-41 | **** |
| 0% NL vs 25% DO | 5.898e-15 | **** |
| 5% DO vs 25% DO | 0.002516 | ** |
| 5% CE vs 25% CE | 0.994 | n.s. |

**Table S3.** Pairwise comparison *p*-values for monolayer, bilayer, and sorting coefficient measurements across phospholipid conditions for TO DEV (**Fig. 5A, B**).

| **BILAYER vs BILAYER comparisons:** |  |  |
| --- | --- | --- |
| Pairs | *P* values | Summary |
| PC vs PC:PE | 0.835 | n.s. |
| PC vs PC: PE: PI | 0.471 | n.s. |
| PC vs PC: PE: PS | 0.461 | n.s. |
| PC: PE vs PC: PE: PI | 0.125 | n.s. |
| PC: PE vs PC: PE: PS | 0.363 | n.s. |
| PC: PE: PI vs PC: PE: PS | 0.002 | ** |
| **MONOLAYER vs MONOLAYER comparisons:** |  |  |
| Pairs | *P* values | Summary |
| PC vs PC:PE | 9.475e-05 | **** |
| PC vs PC: PE: PI | 4.528e-06 | **** |
| PC vs PC: PE: PS | 5.990e-05 | **** |
| PC: PE vs PC: PE: PI | 0.864 | n.s. |
| PC: PE vs PC: PE: PS | 0.678 | n.s. |
| PC: PE: PI vs PC: PE: PS | 0.491 | n.s. |
| **SORTING PARTITION COEFFICIENT comparisons:** |  |  |
| Pairs | *P* values | Summary |
| PC vs PC:PE | 0.583 | n.s. |
| PC vs PC: PE: PI | 0.643 | n.s. |
| PC vs PC: PE: PS | 0.941 | n.s. |
| PC: PE vs PC: PE: PI | 0.446 | n.s. |
| PC: PE vs PC: PE: PS | 0.657 | n.s. |
| PC: PE: PI vs PC: PE: PS | 0.733 | n.s. |

**Table S4.** Pairwise comparison *p*-values for monolayer, bilayer, and sorting coefficient measurements across phospholipid conditions for 50:50 v/v TO:DO DEV (**Fig. 5C, D**).

| **BILAYER vs BILAYER comparisons:** |  |  |
| --- | --- | --- |
| Pairs | *P* values | Summary |
| PC vs PC:PE | 0.773 | n.s. |
| PC vs PC: PE: PI | 0.601 | n.s. |
| PC vs PC: PE: PS | 0.169 | n.s. |
| PC: PE vs PC: PE: PI | 0.769 | n.s. |
| PC: PE vs PC: PE: PS | 0.249 | n.s. |
| PC: PE: PI vs PC: PE: PS | 0.487 | n.s. |
| **MONOLAYER vs MONOLAYER comparisons:** |  |  |
| Pairs | *P* values | Summary |
| PC vs PC:PE | 0.269 | n.s. |
| PC vs PC: PE: PI | 0.035 | * |
| PC vs PC: PE: PS | 0.164 | n.s. |
| PC: PE vs PC: PE: PI | 0.161 | n.s. |
| PC: PE vs PC: PE: PS | 0.676 | n.s. |
| PC: PE: PI vs PC: PE: PS | 0.311 | n.s. |
| **SORTING PARTITION COEFFICIENT comparisons:** |  |  |
| Pairs | *P* values | Summary |
| PC vs PC:PE | 5.938e-03 | ** |
| PC vs PC: PE: PI | 0.074 | n.s. |
| PC vs PC: PE: PS | 0.115 | n.s. |
| PC: PE vs PC: PE: PI | 3.866e-02 | * |
| PC: PE vs PC: PE: PS | 0.053 | n.s. |
| PC: PE: PI vs PC: PE: PS | 0.821 | n.s. |

